# Integrating unmatched scRNA-seq and scATAC-seq data and learning cross-modality relationship simultaneously

**DOI:** 10.1101/2021.04.16.440230

**Authors:** Ziqi Zhang, Chengkai Yang, Xiuwei Zhang

**Affiliations:** Dept of Computational Science and Engineering, Georgia Institute of Technology, Atlanta, GA 30332

## Abstract

It is a challenging task to integrate scRNA-seq and scATAC-seq data obtained from different batches. Existing methods tend to use a pre-defined gene activity matrix (GAM) to convert the scATAC-seq data into scRNA-seq data. The pre-defined GAM is often of low quality and does not reflect the dataset-specific relationship between the two data modalities. We propose scDART (**s**ingle **c**ell **D**eep learning model for **A**TAC-seq and **R**NA-seq **T**rajectory), a deep learning framework that integrates scRNA-seq and scATAC-seq data and learns cross-modalities relationships simultaneously. Specifically, the design of scDART allows it to preserve cell trajectories in continuous cell populations and can be applied to trajectory inference on integrated data.

## Introduction

The availability of single cell multi-modality data provides a comprehensive view of every single cell. Single cell RNA-Sequencing (scRNA-seq) and single cell ATAC-Sequencing (scATAC-seq) respectively measure the gene expression and chromatin accessibility profiles of cells, each being considered as an important aspect of a cell. Recently, techniques which can measure both gene expression and chromatin accessibility in the same cells have been proposed [1, 2, 3], but these technologies are still not widely used, and they can suffer from low sensitivity of one of the data modalities. To make use of the enormous amount of existing data, computational methods have been proposed to integrate scRNA-seq and scATAC-seq data obtained separately for the same cell types in different batches [4, 5, 6, 7], with the aim of building larger datasets and potentially learning the relationship between chromatin region and genes. Following a recent review paper on single cell data integration methods [8], we term scRNA-seq and scATAC-seq data that are not jointly profiled in the same cells as *unmatched data*, and integrating such datasets as the *diagonal integration task*. A diagonal integration method is expected to learn an integrated dataset in the form of either low-dimensional latent embedding or a high-dimensional integrated count matrix, where batch effects are removed and cell identity (e.g. the cluster membership) is preserved from the single-modality dataset to the integrated dataset.

A growing number of computational tools have been proposed for diagonal integration. Some methods aim to learn cell latent embedding such that the latent embedding can be used to reconstruct the original dataset [5, 6, 9]. Some use manifold alignment [10] and aim to learn cell latent embedding by enforcing the latent embedding to preserve the pairwise distances of cells in the original feature space (e.g. gene expression space, chromatin accessibility space, etc) [11, 12]. `Seurat` (v3) [4] maps a query dataset to a reference dataset using canonical correlation analysis, and obtains a new data matrix for the query dataset based on the reference dataset. Most of these methods integrate unmatched scRNA-seq and scATAC-seq datasets into the latent space that preserves the cluster structure in the original datasets, but they do not specifically accommodate the case where the cells form continuous trajectories instead of discrete clusters. When the cells form continuous trajectories instead of discrete clusters, the identity of a cell is the location of the cell along the trajectory. For example, if the trajectory has a structure of a rooted tree, the identity of a cell is reflected by both its branch membership and its pseudotime. Since each cell has a unique branch membership and pseudotime, preserving the cell’s identity in a continuous population can be a more challenging task compared to that in discrete populations with clusters, where multiple cells share the same cluster identity.

On the other hand, a majority of the existing diagonal integration methods [4, 5, 6] require a pre-defined *gene activity matrix* (GAM, also called a region-gene association matrix), representing which genomic regions regulate the expression of which genes, to transform the scATAC-seq data into scRNA-seq data by multiplying the GAM to the scATAC-seq data matrix. The limitations of such practice are: (1) A common way to obtain the GAM is to consider the relative locations between the regions and the gene bodies on the genome and assume that the regulating relationship exists only between regions and genes that are closely located [4, 5, 13]. However, such GAMs can be highly inaccurate as closely located regions and genes do not necessarily have regulatory relationships. (2) Simply multiplying the GAM to the scATAC-seq data to obtain scRNA-seq data makes an assumption of linear relationships between regions and genes in cells, which is often not true in biological systems.

Hereby we propose `scDART` (**s**ingle **c**ell **D**eep learning model for **A**TAC-Seq and **R**NA-Seq **T**rajectory integration), a scalable deep learning framework that embeds data modalities into a shared low-dimensional latent space that preserves cell trajectory structures in the original datasets. `scDART` is a diagonal integration method for unmatched scRNA-seq and scATAC-seq data, which is considered a more challenging task than other integration tasks [8]. It incorporates a neural network which encodes the nonlinear gene activity function that represents the relationships between chromatin regions and genes, named the *gene activity module*. `scDART` allows one to learn the latent space representations of the integrated data and the gene activity module at the same time. It can also take advantage of partial cell matching information as prior: that is, if we know certain cells in the scRNA-seq data should be matched with certain cells in the scATAC-seq data, `scDART` can use this information to obtain improved integration, and we name this version of `scDART` as `scDART-anchor`. `scDART` can also be adapted to any two data modalities which have cross-modality interactions. Even though it is designed for cells that form continuous trajectories, it also works for cells that form discrete clusters.

We have tested `scDART` on three real datasets: two unmatched datasets where the scRNA-seq and scATAC-seq data were not jointly profiled, and a matched dataset where both chromatin accessibility and gene-expression are measured simultaneously in the same cells. We also proposed a simulation procedure to simulate scRNA-seq and scATAC-seq data in the same cells and tested `scDART` on simulated datasets. The simulated datasets allow us to quantitatively evaluate the quality of integration and the learned gene activity function. We have compared `scDART` with existing diagonal integration methods including `LIGER` [5], `Seurat v3` [4], `UnionCom` [12], and `MMD-MA` [11] on both real and simulated datasets. The results show that `scDART` learns a joint latent space for both data modalities that well preserve the cell developmental trajectories, and gene activity function encoding the relationship between chromatin regions and genes that is more accurate than current standard practice.

## Results

### Overview of scDART

The schematics of `scDART` are shown in Fig. 1a. The input of `scDART` are: a scRNA-seq count matrix, a scATAC-seq data matrix, and a pre-defined GAM. The pre-defined GAM is obtained with a commonly used procedure based on genomic locations (Methods) and serves as prior information for `scDART` to learn the gene activity function that more accurately represents the relationship between scATAC-seq and scRNA-seq data. The pre-defined GAM is a binary matrix with rows corresponding to regions and columns corresponding to genes.

**Figure 1:**
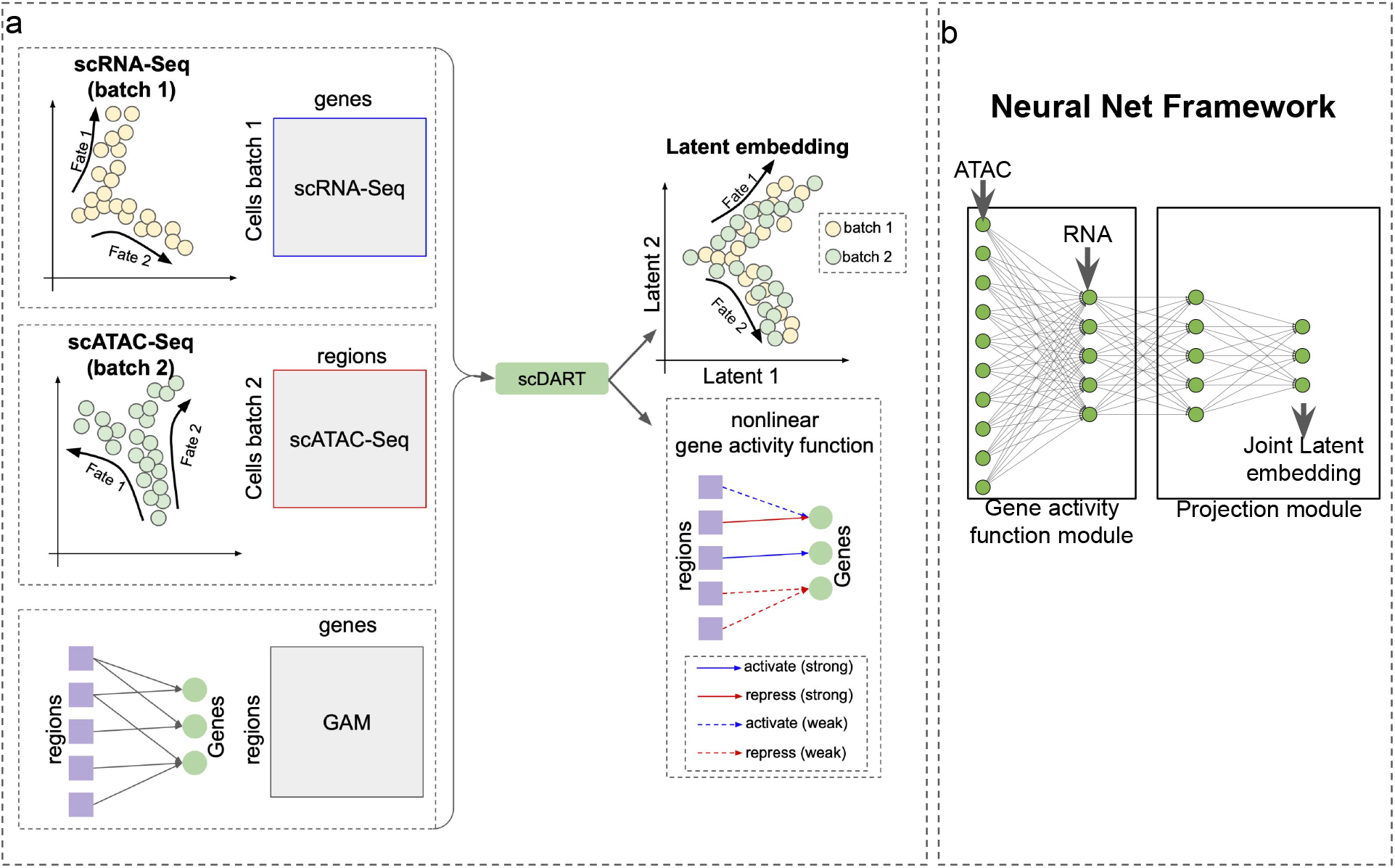
Overview of `scDART`. **a** `scDART` takes as input a scRNA-seq data batch, a scATAC-seq data batch and a pre-defined GAM. It learns the latent embedding of integrated data from the two data batches and a more accurate gene activity function between regions and genes. This gene activity function can be used to predict scRNA-seq data from scATAC-seq data (the predicted scRNA-seq data is also called pseudo-scRNA-seq data). **b** The neural network structure of `scDART`. `scDART` includes two modules: (1) the gene activity function module is a fully-connected neural network. This module encodes the nonlinear regulatory relationship between regions and genes, and generate the pseudo-scRNA-seq data from scATAC-seq data. (2) the projection module takes in the scRNA-seq data and the pseudo-scRNA-seq data, and generates the latent embedding of both modalities.

The neural network structure of `scDART` is shown in Fig. 1b. `scDART` consists of two modules: gene activity function module and projection module. The gene activity function module is a neural network that encodes the nonlinear gene activity function. It takes in the scATAC-seq matrix, transforms the chromatin accessibility of cells into gene expression, and generates a “pseudoscRNA-seq” count matrix. The projection module takes in both scRNA-seq count matrix and the pseudo-scRNA-seq count matrix, and projects them into a shared latent space.

The objective of `scDART` is designed considering three constraints. (1) To preserve cell identity and the trajectory structure in the latent space, `scDART` forces the pairwise Euclidean distances between cells in the latent space to approximate their relative distance along the trajectory manifold in the original feature space (gene expression and chromatin accessibility space). `scDART` uses *diffusion distance* to calculate cell relative distance on the trajectory manifold. *Diffusion distance* has been successfully used by trajectory inference methods [14, 15] to measure the differences between cells along the trajectory. It is advantageous in preserving the trajectory structure in the latent space compared to other distance metrics such as Euclidean or shortest-path distance [16]. In addition, *diffusion distance* can also be directly translated into the pseudotime of cells [14], which facilitates downstream analysis using integrated datasets such as trajectory inference and differential expression (DE) analysis. (2) We consider the scenario where cells in the two batches are sequenced from the same cell types, thus they should have the same trajectory structure. `scDART` uses Maximum Mean Discrepancy (MMD) [17] to measure the similarity of the trajectory structures between the latent embedding of two cell batches, and minimizes the MMD loss such that the cells in two batches “merge” into the same trajectory. (3) `scDART` considers the pre-defined GAM as prior information to assist it to learn the gene activity function module which encodes a more accurate cross-modality relationship than the pre-defined GAM. A novel loss function is designed for `scDART` to incorporate this information.

We design the overall loss function considering all three constraints above. We denote the data matrix of scRNA-seq and scATAC-seq batches respectively as **X**_RNA_ and **X**_ATAC_, the latent embedding of scRNA-seq and scATAC-seq batches as **Z**_RNA_ and **Z**_ATAC_, and the parameters in the gene activity function module and projection module as **Θ**_gact_ and **Θ**_proj_. Then the overall loss function can be written as Equ. 1.

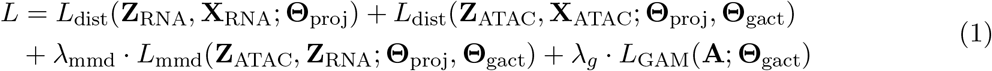

The first two loss terms, *L*_dist_(**Z**_RNA_, **X**_RNA_; **Θ**_proj_) and *L*_dist_(**Z**_ATAC_, **X**_ATAC_; **Θ**_proj_, **Θ**_gact_), measure how well the pairwise Euclidean distances between cells in the latent embedding approximate the diffusion distances between cells, respectively in the scRNA-seq and the scATAC-seq batches. The third term, *L*_mmd_(**Z**_ATAC_, **Z**_RNA_) measures how well the latent embedding of two modalities are “merged” together into the same trajectory structure. The last term allows the gene activity module to incorporate the useful information in the pre-defined GAM. *λ*_mmd_ and *λ*_*g*_ are hyperparameters that control the strengths of the third and forth loss terms. A detailed explanation of each loss term is included in “Methods”.

In certain cases, we have prior information on which cells from the two batches should have the same identity and should be aligned together. For example, the root cells of the trajectory in each batch of data are sometimes known in advance. `scDART` is able to use this information to obtain a better integration. When merging the two cell batches in the latent space, `scDART` takes in the root cells as the anchor cells, and forces the anchor cells in two data batches to merge. The anchor-merge is achieved by adding an anchor loss term *L*_anchor_(**Z**_ATAC_, **Z**_RNA_; **Θ**_proj_, **Θ**_gact_) into Equ. 1. We term the version of `scDART` that uses the anchor cells as `scDART-anchor`. The detailed formulation of *L*_anchor_(**Z**_ATAC_, **Z**_RNA_; **Θ**_proj_, **Θ**_gact_) is included in “Methods”.

The training process of `scDART` is in Methods. After having trained the model and obtained the latent embedding **Z**_ATAC_ and **Z**_RNA_, we apply a post-processing step to further refine the latent embedding to form a cleaner trajectory structure (see “Methods” for more details). The learned cell embedding can be used for trajectory inference and other downstream analyses. In this manuscript, we use Leiden clustering and minimum spanning tree (MST) to detect the trajectory backbones and DPT [14] to infer cell pseudotime (Methods).

### scDART reconstructs cell trajectory and cell matching information in the mouse neonatal brain cortex dataset

To evaluate the performance of `scDART`, we tested it on a mouse neonatal brain cortex dataset obtained by SNARE-Seq [1], where the chromatin accessibility and gene expression profiles were jointly measured for every single cell. The dataset measured 1469 cells on the differentiation trajectory from intermediate progenitors to upper-layer excitatory neurons. We ran `scDART` assuming that chromatin accessibility and gene expression profiles are measured separately from two different cell batches, and evaluated how well `scDART` reconstructs the matching information between cells from the two batches.

We first visualized the latent space embedding of the integrated data learned with `scDART` and baseline methods (Fig. 2a,b for `scDART`, `Seurat`, `LIGER` and `UnionCom` results; Additional file 1: Figure S1a for `MMD-MA` results). Fig. 2a shows the learned latent embedding of different methods, colored by cell types as annotated in the original paper [1]. In these plots, the expected cell trajectory which is a linear trajectory going through “IP-Hmg2”→”IP-Gadd45g”→”IP-Eomes”→”Ex23-Cntn2”→”Ex23-Cux1” should be preserved. To test this, we took the centroid of cells in each cell type (large red dots in the plots) and applied MST on these points using Euclidean distance between the centroids to obtain the trajectory backbone (red lines in plots). One can observe that only `scDART` clearly shows the expected trajectory. Fig. 2b visualizes the integrated latent embedding colored with batch (or modality). In these plots, one expects to see that the two batches are merged and mixed. All methods merge the two batches except for `Seurat`.

**Figure 2:**
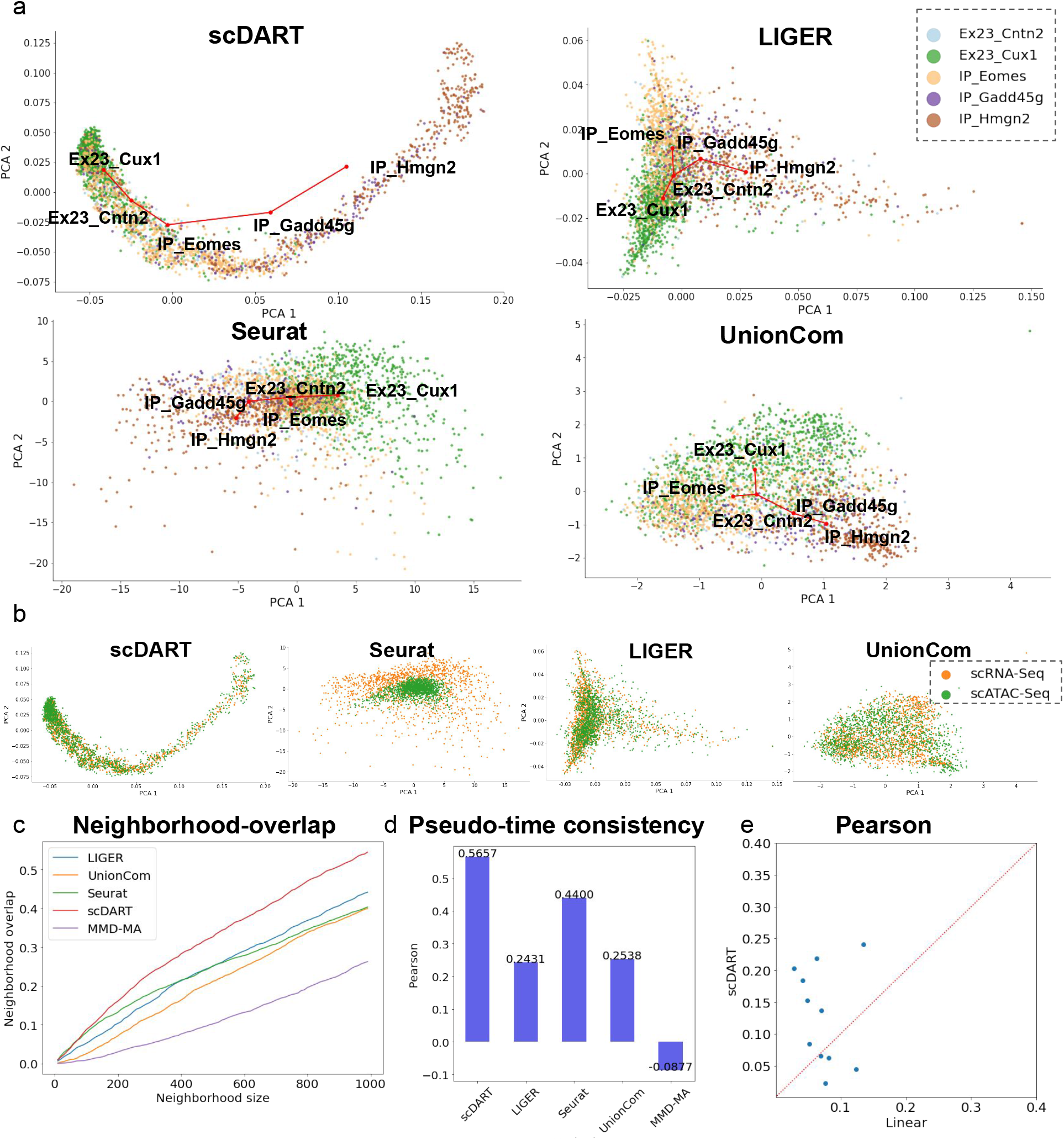
The results of `scDART` and baseline methods on the SNARE-seq mouse neonatal brain cortex dataset. **a** Latent embedding of `scDART`, `LIGER`, `Seurat`, and `UnionCom`, visualized using PCA. Cells are colored with cell type annotation in the original paper. Red lines show the inferred trajectory backbone. All plots share the same legend as in the `LIGER` plot. **b** Latent embedding of `scDART`, `Seurat`, `LIGER`, and `UnionCom`, where cells are colored with data batches. All plots share the same legend as in the `UnionCom` plot. **c** Neighborhood overlap score. **d** Pseudotime consistency score, where Pearson correlation is used. **e** Pearson correlation between pseudo-scRNA-seq (predicted scRNA-seq from scATAC-seq data), learned by `scDART` (y-axis) and linear transformation (x-axis), and real-scRNA-seq. Top 20 DE genes are shown.

Based on the predicted trajectory and cell pseudotime from `scDART`, we analyzed the differentially expressed (DE) genes and motifs along the trajectory. Using the latent embedding of `scDART`, we inferred cell pseudotime on each modality separately, and found genes that are DE with pseudotime using likelihood ratio test (see “Methods” for more details). Out of the DE genes we found, there are *Mki67* and *Fabp7* that are expected to be highly expressed in the initial stage of the trajectory [1], *Eomes* and *Unc5d* that are abundant in the neuroblast stages (their expression first increases then decreases), and *Cux1* and *Foxp1* that mark the upper-layer neurons (Additional file 1: Figure S1b). These findings are consistent with the original paper [1].

We then transformed the chromatin accessibility into motif deviation using `ChromVAR`, and detected the differentially accessible motifs along the trajectory (see “Methods” for more details). We found motif of transcription factor *Neurog1* (“MA0623.1_Neurog1”), a common regulator that involves in the initialization of neuron differentiation. We also found motif of transcription factors of SOX family such as *SOX6* (“MA0515.1_Sox6”), *SOX10* (“MA0442.1_SOX10”), *SOX2* (“MA0143.3_Sox2”), etc, which regulates the nervous system development (information from GeneCards [18]). The full lists of differentially expressed genes and accessible motifs are available in Additional file 2: supplementary table 1.

Since this dataset has ground-truth cell matching information, we can quantify how well `scDART` integrates the two modalities in addition to the visualization in Figs. 2a,b, and compare it with baseline methods including `Seurat`, `LIGER`, `UnionCom`, and `MMD-MA`. We first calculate the *neighborhood overlap score*, which measures how many matched cells are located in the close neighborhood of each other in the latent space (see “Methods” for more details). The matched cells should be embedded into the exact same location as they have the same original cell identity. We calculate the neighborhood overlap score for different neighborhood size, and plot the curve in Fig. 2c. The curve shows that `scDART` recovers cell matching better compared to the other baseline methods. In addition, we also quantify the consistency of pseudotime inferred from cells in each modality, since matched cells should ideally have the same pseudotime along the trajectory. We first infer the pseudotime of each data modality on the latent space separately, then calculate the correlation of pseudotime between matched cells using Pearson and Spearman correlation. The result shows that `scDART` has the best consistency and `Seurat` also achieves good performance close to `scDART` (Fig. 2d, Additional file 1: Figure S1c).

Finally, we can also use this jointly profiled dataset to test the non-linear gene activity function learned by `scDART`. Since for each cell in the dataset that is measured with scATAC-seq, we also know its corresponding scRNA-seq profile. We then compare the real scRNA-seq profile and the “pseudo-scRNA-seq” predicted from the gene activity module using the scATAC-seq data as input. We specifically investigated the DE genes along the pseudotime inferred with the gene expression modality. First, we visualize the predicted “pseudo-scRNA-seq” data and the real scRNA-seq data on these genes with heatmap (Additional file 1: Figure S2a), and observe similar changing patterns of genes between the two plots.

We then compare the gene activity function of `scDART` with a linear transformation on the chromatin accessibility data using the input GAM in terms of the ability of predicting gene expression data from chromatin accessibility data. We quantify this ability by calculating Pearson correlation between predicted gene expression data and measured gene expression data. From Additional file 1: Figures S2b,c, we can observe that for most of the genes the correlation scores obtained by `scDART` are higher.

### scDART integrates mouse endothelial cell development datasets

We tested `scDART` on the mouse endothelial cell development dataset from [19]. The authors conducted scATAC-seq and scRNA-seq separately on two batches of mouse endothelial cells, where scATAC-seq measured one batch of 1186 cells and scRNA-seq measured another batch of 1628 cells. The cells undergo a differentiation path from endothelial cells (Endo) to Hematopoietic stem and progenitor cells (HSPCs) that accumulate in intra-arterial clusters (IAC). The overall cell trajectory in this dataset is a linear-like path, where endothelial cells differentiate into “Arterial Endo 1/2” and then become “Pre-BN1” cells. “Pre-BN1” cells then undergo a maturation path to “IAC” through “Pre-HE” and “HE” stages.

We obtained the latent embedding of the integrated dataset using `scDART` and visualized it in Figs. 3a-c. We then applied our trajectory inference procedure (Methods) to the latent embedding to infer the trajectory backbone (Fig. 3a) and cell pseudotime (Fig. 3c). The results (Figs. 3a-c) show that `scDART` is able to integrate scATAC-seq and scRNA-seq batches into the same latent space while preserving the linear trajectory structure. The sudden drop of the density of cells on the trajectory, where the cells are distributed densely at the beginning and then becomes very sparse at the end of the path, corresponds to the developmental “bottleneck” between “Pre-HE” and “HE” discussed in [19]. It also shows that there may be a “speeding up” of cell differentiation towards the end of the trajectory. We also performed baseline methods including `LIGER`, `Seurat`, `UnionCom`, and `MMD-MA` on the datasets, and visualize their latent embedding using PCA (Figs. 3d,e, Additional file 1: Figure S3a). `LIGER` is able to merge the two batches of cells (Fig. 3e, top plot) but does not preserve the linear trajectory (Fig. 3d, top plot). `Seurat` shows a linear trajectory (Fig. 3d, middle plot) but cells from the two batches are not well integrated with the cells from scATAC-seq being more concentrated than those from the scRNA-seq batch (Fig. 3e, middle plot). `UnionCom`, while preserves the linear trajectory, mis-integrates “HE”, “Pre-BN1” and “IAC” cell types (Figs. 3d,e, bottom plots). `MMD-MA` also fails to merge the cells from different batches (Additional file 1: Figure S3a).

**Figure 3:**
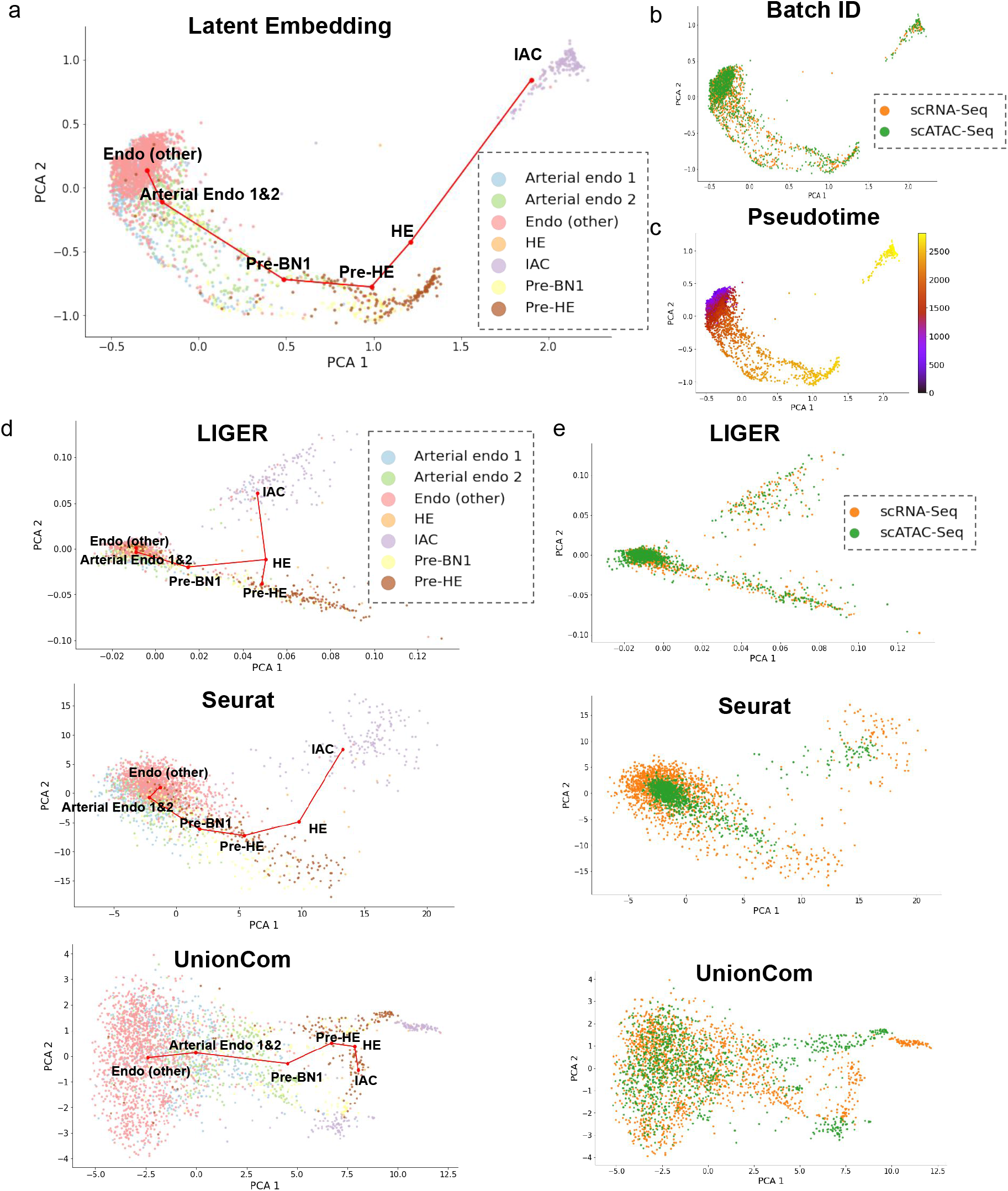
The results of `scDART` and baseline methods on the mouse endothelial cell development dataset. **a-c** Latent embedding of `scDART`, visualized using PCA. Cells are colored with (a) cell type annotation from original data paper, (b) data batches, and (c) inferred pseudotime. Red lines in (a) show the inferred trajectory backbone. **d** The latent embedding of `Seurat`, `LIGER`, and `UnionCom` where cells are colored with cell type annotation from original data paper. All plots share the same legend as in the `LIGER` plot. **e** The latent embedding of `Seurat`, `LIGER`, and `UnionCom` where cells are colored with data batches. All plots share the same legend as in the `LIGER` plot.

We then analyzed the DE genes and differentially accessible motifs along the trajectory (see “Methods” for more details). We further performed gene enrichment analysis using `TopGO` [20] on the DE genes, and found multiple enrichment terms relevant to hematopoietic stem and progenitor cells generation and other embryonic maturation processes. We found “myeloid cell differentiation”, “positive regulation of cell development”, “regulation of stem cell proliferation” in gene enrichment terms (Additional file 1: Figure S3b, the full list is incorporated in Additional file 3: supplementary table 2). Then we analyzed the motifs using `ChromVAR`, and found enriched motifs using likelihood ratio test (see “Methods” for more details). Interestingly, we found multiple motifs of transcription factors including *RUNX1, GATA, SOX*, and *FOX*. Those TF motifs are also reported in the original paper to be closely related to hematopoietic stem and progenitor cells generation process (Additional file 1: Figure S3c).

### scDART integrates human hematopoiesis datasets and learns myeloid and lymphoid cell trajectories

We applied `scDART` to a human hematopoiesis dataset and analyzed the biological factors that drive the differentiation process from hematopoietic stem and progenitor cells (HSC) to myeloid and lymphoid cells. We collected scATAC-seq data from Buenrostro *et al* [21], and scRNA-seq data from Pellin *et al* [22]. 1367 cells from the scATAC-seq batch and 1666 cells from scRNA-seq batch were used. Myeloid and lymphoid cells are originated from HSC. HSC first develop into multipotent progenitors (MPP), then they undergoes two potential differentiation branches until maturity: the Lymphoid-committed branch (CLP branch) and the Erythroid-committed branch (MEP branch). Cells in CLP branch first transit into Lymphoid multipotent progenitors (LMPP) and mature into common lymphoid progenitor cell (CLP), whereas cells in MEP branch undergo a differentiation path to megakaryocyte-erythroid progenitor (MEP) cells through common myeloid progenitor (CMP) cells. Note that in both papers presenting the original datasets [21, 22] the HSC and MPP cells were not separated and following these papers we use “HSC&MPP” to represent both cell types. Therefore, the expected trajectory on this dataset is a bifurcating trajectory with HSC&MPP as root cell type.

Figs. 4a,b show the latent embedding of `scDART`. Cells from different data batches are well integrated by `scDART`. At the same time, the inferred trajectory on the latent embedding (backbone as the red lines in Fig. 4a, pseudotime in Additional file 1: Figure S4a) also shows a clear bifurcating pattern following the differentiation path of HSC.

**Figure 4:**
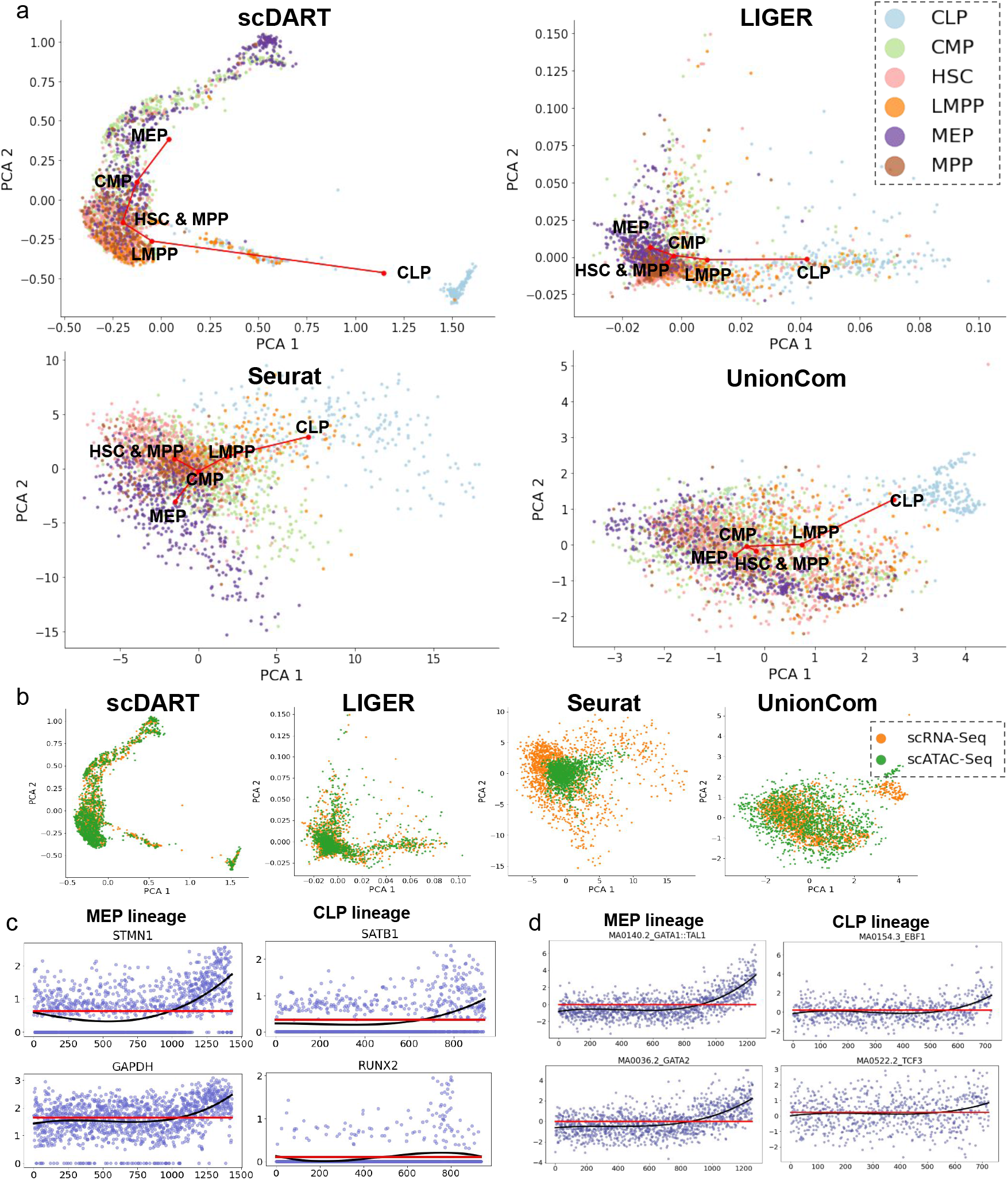
The results of `scDART` and baseline methods on the human hematopoiesis dataset. **a** Latent embedding of `scDART`, `LIGER`, `Seurat`, and `UnionCom`. Cells are colored with cell type annotations from the the original paper. Red lines show the inferred trajectory backbone. All plots share the same legend as in the `LIGER` plot. **b** Latent embedding of `scDART`, `Seurat`, `LIGER` and `UnionCom`. Cells colored with data batches. All plots share the same legend as in the `UnionCom` plot. **c** The expression level of *STMN1* and *GAPDH* along MEP lineage, and *SATB1* and *RUNX2* along CLP lineage. Cells are ordered on the x-axis according to the inferred pseudotime. **d** The deviation values (from ChromVAR) of differentially accessible motifs along MEP and CLP lineages. The black and red lines in (c) and (d) correspond to the fitted statistical models under alternative and null hypothesis, respectively, when conducting likelihood ratio test.

In terms of baseline methods, even though `LIGER` and `UnionCom` can integrate cell batches, their latent embedding fail to show a correct trajectory that distinguishes myeloid and lymphoid cell branches (Figs. 4a,b). Both methods mistakenly assigned CMP as the branching point as the two branches were not sufficiently separated in their latent spaces. `Seurat` and `MMD-MA` both have difficulty integrating cells from the two batches (Figs. 4a,b, Additional file 1: Figure S4b).

We further analyzed the DE genes along both the MEP and CLP branches inferred from scRNAseq data (see “Methods” for more details). We found DE genes such as *STMN1* and *GAPDH* in MEP branch, and *SATB1* and *RUNX2* in CLP branch. These genes was also shown to be the marker genes for the two branches in [22] (Fig. 4c). We conducted gene ontology enrichment analysis using `TopGO`. The top enriched terms (Additional file 1: Figure S4c) include branch specific terms such as myeloid leukocyte differentiation for MEP branch, and B cell receptor signaling pathway for CLP branch. We also find terms that are relevant to general cell differentiation process. We obtained the TF motifs deviation score from scATAC-seq data using `ChromVAR`, and analyzed the differentially accessible motifs along both branches (see “Methods” for more details). We found motifs of GATA TF class (‘MA0140.2_GATA1::TAL1’, ‘MA0036.2_GATA2’, ‘MA0037.2_GATA3’, left column in Fig. 4d and top figure in Additional file 1: Figure S4d) along MEP branch, which is the key regulator of MEP branch [21]. We also found motifs related to TFs such as EBP1 (‘MA0154.3_EPB1’), TCF3 (‘MA0522.2_TCF3’), TCF4 (‘MAO830_TCF4’) along the CLP branch (right column in Fig. 4d and bottom figure in Additional file 1: Figure S4d). Those TFs were also reported in [21] as the key regulators of the CLP branch.

### Testing scDART using simulated data

We proposed a simulation procedure which allows us to simulate scRNA-seq and scATAC-seq data in the same cells. Our simulation process considers that the chromatin accessibility data affects the probability that a gene’s expression is switched *on* or *off*, and jointly simulates scRNA-seq and scATAC-seq data with this relationship (Additional file 1: Figure S5a, Methods). We simulated multiple scRNA-seq and scATAC-seq datasets with different trajectory topology, as well as dataset with discrete clusters. Each dataset can have two or more batches, where both modalities are simulated for cells in every batch. From the simulated matched data, we can obtain unmatched data by keeping only one modality in one batch to test diagonal integration methods.

### Quantifying latent embedding accuracy on simulated datasets

Using simulated data with continuous trajectories, we quantify the performance of `scDART` and `scDART-anchor`, along with baseline methods including `LIGER`, `Seurat`, and `UnionCom`.

We first tested how well the cell latent embedding preserves the original trajectory structure. We simulated 6 datasets with different trajectory topologies: 3 bifurcating trajectory topology and 3 with trifurcating trajectory topology. The backbones of different trajectory structures used in simulation are shown in Fig. 5a. Using simulated data, we ran the diagonal integration methods and inferred trajectories on the latent embedding of each method (see “Methods” for details). Then, we measured the accuracy of cell branch assignment using F1 score [23]. We also measured how well the inferred pseudotime matches the true pseudotime along the trajectory using Kendall-*τ* score (also known as Kendall rank correlation). More details on F1 and Kendall-*τ* scores are in Methods. Since `scDART` and `scDART-anchor` include randomness when sampling the mini-batch in stochastic gradient descent, we ran them with 3 different random seeds for each dataset. When running `scDART-anchor`, we use 10 root cells with the smallest pseudotime from each dataset as the anchor.

**Figure 5:**
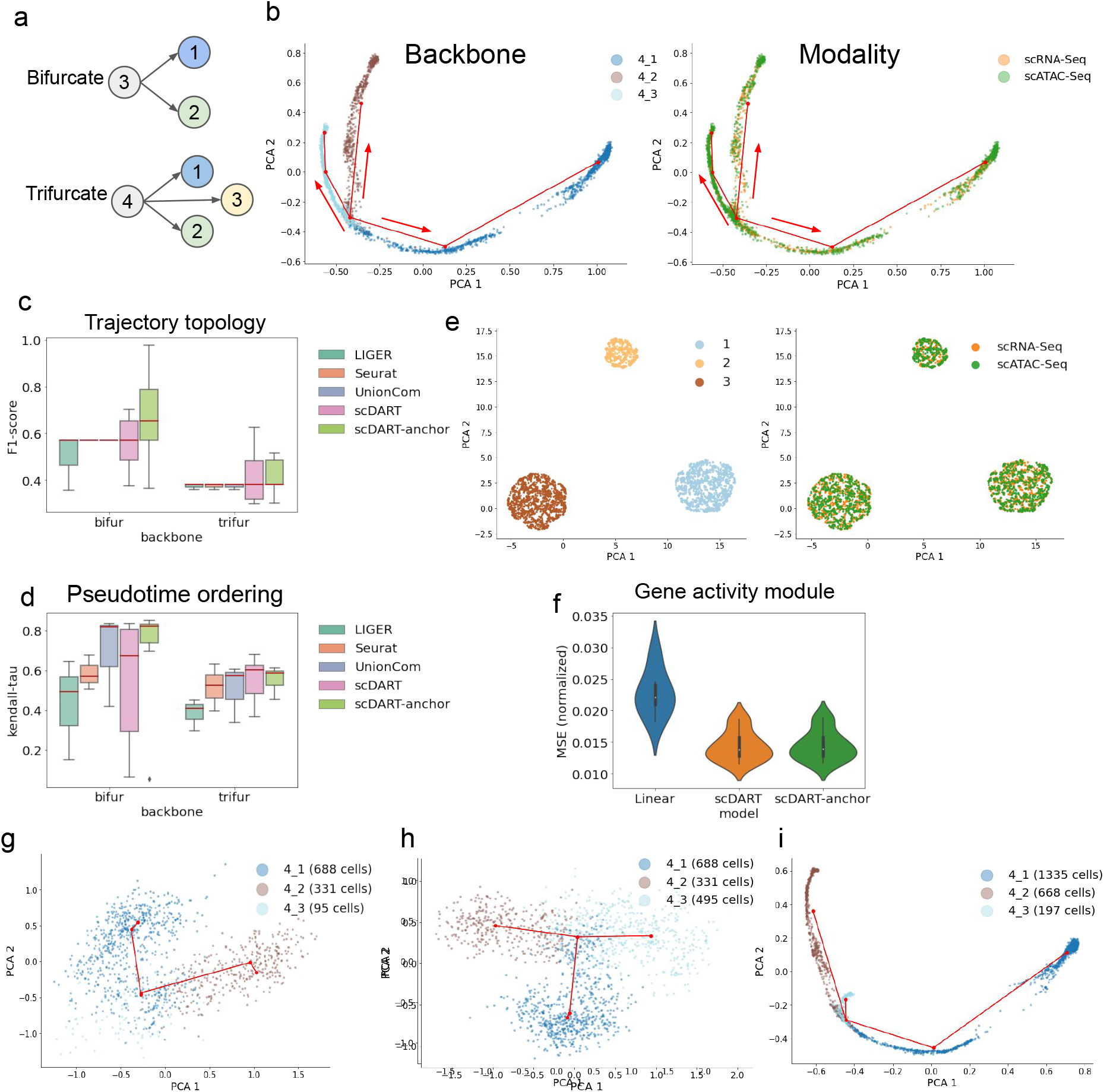
Performance of `scDART` on simulated datasets. **a** Ground truth trajectory topology of simulated continuous datasets. **b** The latent embedding of `scDART` from one simulated data with trifurcating trajectory backbone. Red lines show the inferred backbone, and arrows show the trajectory direction. Left: cells colored by the ground truth trajectory branches they belong to; Right: cells colored by the data batches. **c-d** Boxplotx of F1-score and Kendall-*τ* score calculated from different methods. 3 datasets were used for each type of trajectory, and for each dataset `scDART` and `scDART-anchor` were run 3 times. `Seurat` and `UnionCom` have the same F1 score on all 3 datasets with bifurcating trajectory. **e** The latent embedding of `scDART` on one dataset with discrete clusters. Left: cells colored with cell type annotations; Right: cells colored with data batches. **f** Violin plots of normalized MSE between pseudo-scRNA-seq and ground truth scRNA-seq. **g** Trajectory backbone learned from scRNA-seq when branch 4 3 has only 95 cells. **h** Trajectory backbone learned from scRNA-seq when branch 4 3 has 495 cells. **i** Trajectory backbone learned from latent embedding integrated by `scDART` where when branch 4 3 has 197 cells (95 from scRNA-seq and 102 from scATAC-seq).

One sample result of scDART is visualized in Fig. 5b. `scDART` is able to integrate the cells from two data batches into the latent space where the trifurcating trajectory pattern is preserved. The boxplots of scores are shown in Figs. 5c,d. Fig. 5c shows that `scDART` learns latent embedding that is able to well preserve the trajectory topologies, especially when the root cell information is provided. `LIGER` has the lowest F1 score mainly because the latent embedding of cells is not stretched apart enough along the cell pseudotime, which makes the trajectory inference algorithm detect the wrong branching structure. In Fig. 5d, `scDART` has some low Kendall-*τ* score for bifurcating topologies. The main reason is that the simulated data has almost the same cell density along the trajectory, which makes `scDART` mistakenly detect the wrong starting and ending point of those simple trajectory topologies. Such an equal-density scenario rarely happens in real dataset, as the cell does not always differentiate at the same speed, and it can be solved by using the anchor cell information. `scDART-anchor` stably achieves much higher F1 and Kendall-*τ* scores than other methods when using the root cell information.

We also use the simulated datasets to show that `scDART` can not only integrate data from continuous populations but also integrate data from discrete clusters. We generated simulated datasets with discrete cluster structure (3 clusters, scRNA-seq batch has 1530 cells and scATAC-seq batch has 1470 cells), and used `scDART` to learn the latent embedding. The results (Fig. 5e) show that `scDART`, while being able to preserve continuous trajectories during integration, can be applied to discrete clusters too.

### Test gene activity module accuracy on simulated datasets

We use the gene activity module in `scDART` to encode the data-specific regulatory relationship between regions and genes, which meanwhile helps the model to learn a better cell embedding. We validate the capability of predicting scRNA-seq data from scATAC-seq data of this module on simulated data, and compare the results with baseline procedure where the input GAM to `scDART` is used to for the prediction by linearly transforming the scATAC-seq data matrix into the scRNA-seq data matrix. Normalized mean square error between the predicted scRNA-seq data (also called pseudo-scRNA-seq data) and the ground truth scRNA-seq data was used as evaluation metric (see “Methods” for more details).

To prepare the data used for this test, we first generated *N* cells with jointly profiled scATAC-seq and scRNA-seq data using our simulation procedure, then divided these *N* cells into 2 batches with respectively *N*_1_ and *N*_2_ cells, and used the scRNA-seq data from batch 1 and the scATAC-seq data from batch 2 as the input for `scDART`. We take the output data from the gene activity module as the corresponding pseudo-scRNA-seq data for scATAC-seq data from batch 2, and compare it with the ground truth scRNA-seq data of batch 2.

The baseline method is to directly multiply GAM with the scATAC-seq data to obtain the pseudo-scRNA-seq data from batch 2, as is used in existing integration methods including `Seurat` and `LIGER`. Note that with our simulated procedure we have true GAMs with binary values and we used the true GAMs for the baseline method. We termed the baseline method as *linear prediction*. We generated 6 simulated datasets for the test. The resulting violin plot (Fig. 5f) shows that `scDART` has a much lower error compared to *linear prediction* even when the true GAM was used in the *linear prediction*. `scDART-anchor` has a similar performance as `scDART`.

### scDART detects branches with small number of cells

Performing integration of single cell data from more than one modality is expected to reveal knowledge that can not be learned with single-modality data. On datasets where cells form discrete clusters, some methods were shown that they can detect rare cell types on the integrated dataset [4, 5, 24]. In the case of continuous trajectories, a branch in a trajectory may not be detected if the number of cells on that branch is very small in a dataset. Integrating this dataset with other datasets may help recover this branch. We show that scDART can detect such branch after integration.

Using our simulation procedure we simulated one batch of scRNA-seq data and one batch of scATAC-seq data with a trifurcating ground truth trajectory where one branch is shorter than others and cells are also sparse on this branch (PCA visualization of scRNA-seq in Fig. 5g, and scATAC-seq in Additional file 1: Figure S5b, where the small branch is “4_3”). Branch “4_3” cannot be detected when we apply the trajectory inference procedure (Methods) on the scRNA-seq dataset after its dimension is reduced by PCA.(Fig. 5g). The branch cannot be detected with only scATAC-seq dataset either (Additional file 1: Figure S5b, with the scATAC-seq data we used latent semantic indexing (LSI) for dimensionality reduction instead of PCA following [25, 26]). It is possible that applying a different dimensionality reduction method to the scRNA-seq data may allow branch “4_3” to be detected, but we show that the small number of cells on this branch is the main cause for the branch being undetected, by the results in Fig. 5h: the difference between the scRNA-seq datasets used in Fig. 5g and Fig. 5h lies in the number of cells on branch “4_3”. For the former there are 95 cells and for the latter there are 495 cells. Using the same trajectory inference procedure, branch “4_3” can be detected in Fig. 5h. And same for the case of scATAC-seq data (Additional file 1: Figures S5b,c). After integrating the scRNA-seq and scATAC-seq data sets (scRNA-seq has 95 cells on the branch “4_3” and scATAC-seq has 107 cells on the branch “4_3”) using scDART, branch “4_3” can be easily detected with the simple trajectory inference procedure (Fig. 5i and Additional file 1: Figure S5d), which shows the capability of `scDART` in detecting small branches.

### Effects of hyper-parameters

The hyper-parameters in `scDART` (`scDART-anchor`) include the latent dimension *k* and the weights of loss terms in the loss function (Equ. 1), *λ*_*g*_ and *λ*_*mmd*_. The latent dimension *d* is determined by the complexity of the trajectory structure in the data. A larger *d* is needed for datasets with complex trajectory structures. We used *d* = 4 for all real datasets and *d* = 8 for simulated continuous datasets as simulated datasets include bifurcating and trifurcating structures. *λ*_*g*_ controls how strong the prior gene activity matrix affects the training of the gene activity module. In all results we presented in this manuscript, we used the default value *λ*_*g*_ = 1. *λ*_*g*_ can be adjusted according to the quality of the input GAM — larger *λ*_*g*_ can be used if users have high confidence in the input GAM. *λ*_*mmd*_ controls how well the latent distributions of cells in different batches are “merged”. In most of our test results we also used the default value *λ* = 1, and only on the mouseneonatal brain cortex dataset we set *λ* = 10 for stronger merging effect.

We comprehensively tested the robustness of `scDART` and `scDART-anchor` against different hyper-parameter settings using 4 simulated datasets selected from the datasets used in the previous tests. The datasets include both the bifurcating and trifurcating trajectories. For each dataset, we measured the F1 score and Kendall-*τ* score of our methods under different combinations of hyper-parameter values, where the values of each parameter are as follows: *λ*_*g*_ = 0.1, 1, 10, *λ*_*mmd*_ = 1, 10, and *d* = 4, 8, 32. For each dataset under each hyper-parameter setting, we run `scDART` and `scDART-anchor` with 3 different random seeds. The results are summarized as boxplots in Additional file 1: Figures S5e,f. The results show that the F1 score lies within a stable range between 0.5 to 0.6, and the median Kendall-*τ* score also stays at around 0.6 under different hyper-parameter settings, which shows a robust performance of the model to hyper-parameters *d, λ*_*mmd*_ and *λ*_*g*_. In addition, the comparison between the boxplots of `scDART` and `scDART-anchor` shows that higher robustness can be achieved when the anchor information is given.

## Discussion

Through `scDART`, we have shown the effectiveness of learning the integrated data and cross-modality relationship simultaneously. Based on this idea, new methods can be developed to address some limitations in the current `scDART` model. First, like most existing methods, `scDART` is not designed for data batches where cells have different trajectories in different batches. Addressing disparities with continuous trajectories can be even more challenging than with discrete clusters, as disparities with continuous populations include various scenarios: additional or missing branches in the trajectories, different lengths of certain branches, different orders of cells on certain branches. We anticipate that the cross-modality relationship will play a more important role in methods integrating data with trajectory disparities.

Although the gene activity module learned in `scDART` can predict scRNA-seq data from scATAC-seq data more accurately compared to the conventional approach of linear transformation with GAM, the gene activity module is not perfect in performing this prediction task. In `scDART`, it has helped the integration task. To learn a highly accurate function that predicts scRNA-seq data from scATAC-seq data, matched (jointly-profiled) datasets can be leveraged for cell types where such data is available.

Currently, `scDART` works with two modalities, chromatin accessibility and gene expression. The framework can be generalized to work with three modalities, for example, chromatin accessibility, gene expression and protein abundance by adding another neural network representing the relationship between gene expression and protein abundance.

## Conclusions

Although technologies which can jointly profile more than one modalities are available, a majority of the existing single-cell datasets are single-modality data and diagonal integration methods are needed to integrate different modalities from different batches. We proposed `scDART`, which is a diagonal integration method for scRNA-seq and scATAC-seq data, with the following advantages compared to existing methods: (1) existing methods use a pre-defined generic GAM to convert the scATAC-seq data into scRNA-seq data or map the manifold of the two data modalities without using the GAM. `scDART` learns the relationship between the scATAC-seq and the scRNA-seq data represented by a neural network, which is data-specific and can be nonlinear; (2) existing diagonal integration methods are heavily tested using datasets where cells form discrete clusters. `scDART` is particularly designed to preserve continuous trajectories in the integrated datasets and this strength of `scDART` has been shown by the comparison between `scDART` and existing methods using continuous populations of cells.

To our knowledge, `scDART` is the first method that performs the two important tasks, integrating two data modalities which are not jointly profiled and learning the cross-modality relationships, simultaneously in the case of scATAC-seq and scRNA-seq data. In the era of single cell multi-omics, the goal of data integration should be not only removing batch effects and compiling larger datasets, but also learning the relationship between different data modalities. We expect that more methods which can learn relationship across modalities will be developed in the future to take advantage of the multi-modal omics data.

## Methods

The detailed loss terms in Equ. 1 are described in sections below. *L*_dist_(·) is described in the section *Preserving trajectory structure in latent embedding* ; *L*_mmd_(·) is described in the section *Integrating modalities and batch removal* ; *L*_gact_(·) is described in the section *Using prior gene activity matrix. L*_anchor_(·) is described in the section *Using anchor information*. In addition to the loss terms, the input GAM construction steps is described in the section *Constructing predefined gene activity matrix (GAM)*. The model training procedure is described in the section *Training scDART*. After training the model, the post-processing step is described in the section *Post-processing after training*. The trajectory inference and differential expression that we used in the analysis above is described in the section *Trajectory inference* and *Differential expression analysis*. Simulated data generation procedure and the evaluation metric are described respectively in section: *Data simulation* and *Evaluation metrics*.

### Constructing pre-defined gene activity matrix (GAM)

`scDART` constructs the pre-defined GAM as a binary matrix with rows corresponding to regions and columns corresponding to genes. `scDART` assumes the regions that lie within 3000 base-pairs upstream of the gene body on the genome to be the regulatory regions of that gene, and assign 1 to the corresponding elements in GAM, 0 to the remaining elements.

### Preserving trajectory structure in latent embedding

The trajectory structure and the relative locations of cells on the trajectory can be represented by their pairwise diffusion distances calculated using their gene expression and chromatin accessibility features [14, 15]. In order to preserve trajectories in the latent space, we aim to minimize the difference between the pairwise distances in the latent space and in the original dimensional space. The pairwise distance between cells in the original space is calculated with diffusion distance and the pairwise distance in the latent space is calculated using Euclidean distance. We use *L*_dist_ to denote the difference between cells’ Euclidean distance on the latent embedding and their diffusion distance and we would like to minimize *L*_dist_. *L*_dist_ is calculated separately for scATAC-seq batch and scRNA-seq batch.

The calculation of the diffusion distance matrix is similar to [15]. Given a data matrix **X** (can be either the scRNA-seq or the scATAC-seq data matrix), we first reduce the feature dimension (using PCA for scRNA-seq and Latent Semantic Indexing for scATAC-seq). Then we construct a pairwise similarity matrix **K** using their dimension-reduced representation. More specifically, the similarity between cells *i* and *j* (corresponding to the (*i, j*)*th* element in **K**) is calculated as:

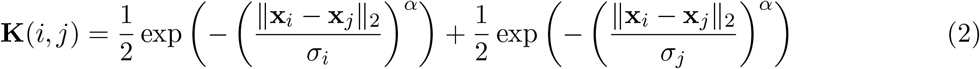

where **x**_*i*_ and **x**_*j*_ are the dimension-reduced representation of cell *i* and cell *j*. The bandwidth *σ*_*i*_ (*σ*_*j*_) is set to be proportional to the distance between cell *i* (*j*) and its *k*th-nearest neighbor (*k* is set to 5). The decay parameter *α* is set to 40 (following default setting used in [15]). The cell transition matrix **P** is constructed by normalizing the similarity matrix **K** such that values in each row sum up to 1: **P** = **K***/*(∑_*j*_ **K**_*ij*_). Then, the diffusion process is performed by powering the transition matrix **P** to *t* times to obtain **P**_*t*_ = **P**^*t*^. The diffusion step *t* is the key parameter in calculating the diffusion distance. Small *t* may not be enough to remove the noise in the dataset; large *t*, on the other hand, may remove noise and useful biological information at the same time. Inspired by [14] who summed up *P*_*t*_ of all *t* values from one to infinity to eliminate *t*s, we calculate the *P*_*t*_ with multiple *t* values (*t* = 30, 50, 70) selected from both small *t*s and large *t*s, and use the averaged 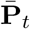 following

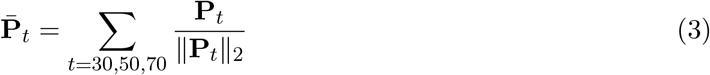

Finally, considering each row of **P**_*t*_ to be the feature vector of the corresponding cell, the diffusion distance matrix **D**_*X*_ can be calculated as the pairwise Euclidean distance between cells in this feature space. After calculating the diffusion distance matrix **D**_*X*_, we then calculate the Euclidean distance between every pair of cells using their latent embedding and obtain distance matrix **D**_*Z*_. *L*_dist_ then measures the difference between **D**_*X*_ and **D**_*Z*_ using KL-divergence:

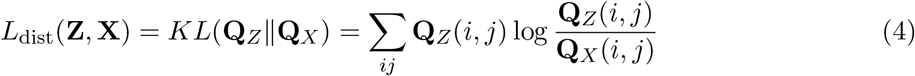

where **Q**_*X*_ and **Q**_*Z*_ are normalized distribution matrix calculated from **D**_*X*_ and **D**_*Z*_:

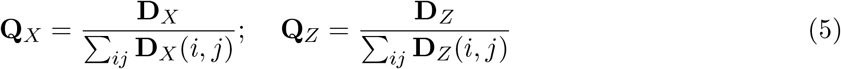

Compared to other possible functions to measure the difference between **D**_*X*_ and **D**_*Z*_, for example, mean square error or inner product loss, the asymmetric formulation of KL-divergence loss (Equ. 4) has a larger penalty when **Q**_*X*_(*i, j*) is small and **Q**_*Z*_(*i, j*) is large, and this will force the latent embedding to better preserve the local manifold structure. This was also discussed in [27].

### Integrating modalities and batch removal

Following existing work [12, 28], we assume the underlying trajectory structures of the input scRNA-seq and scATAC-seq datasets are similar as the cells follow the same biological process. In order to project scRNA-seq data and scATAC-seq data into the same latent space where the trajectory topology of both datasets merge, we incorporate the maximum mean discrepancy (MMD) loss (Equ. 6) [17]. MMD provides a statistical measure of the difference between the distributions of the latent embedding of scRNA-seq and scATAC-seq data. Denoting the latent embedding of scRNA-seq and scATAC-seq data respectively as **Z**_RNA_ and **Z**_ATAC_, the MMD loss function takes the following form:

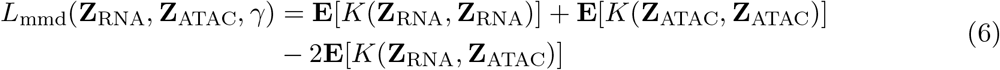

Where **E** means expectation; *K* is a Gaussian kernel function of the form:

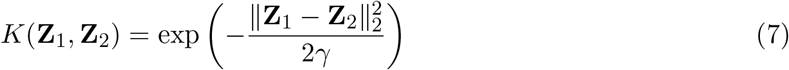

*γ* is a key parameter of the Gaussian kernel function. Following [29], we sum up the MMD loss with different *γ* values to improve the robustness of the loss term:

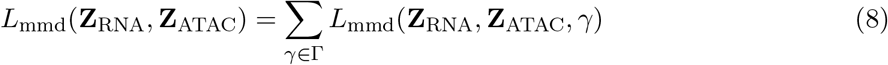

where Γ = *{*10^*u*^*}* and *u* is an integer ranging from −6 to 6.

### Using prior gene activity matrix

We denote the GAM that is obtained from the section *Constructing pre-defined gene activity matrix (GAM)* by **A**. Some existing methods which integrate scRNA-seq and scATAC-seq data multiply **A** to the scATAC-seq data matrix and obtain another scRNA-seq data matrix [5, 4, 6], which is a

linear transformation. However, such transformation is highly inaccurate. How the accessibility of a genomic region affects the expression level of a gene is a complex mechanism, which can be both nonlinear and cell-type specific.

We utilize a three-layer fully connected neural network, termed the “gene activity module”, to learn a gene activity function that can transform scATAC-seq data into scRNA-seq data. The gene activity module thus represents the data-specific relationship between scATAC-seq and scRNA-seq data, and it can encode nonlinearity in this relationship.

The network has an input dimension equal to the number of regions in scATAC-seq data, and an output dimension equal to the number of genes in scRNA-seq data. We use leaky rectified linear unit (ReLU) [30] as the activation function between the layers and remove the bias term of each layer. Taking the region accessibility of a cell (**x**_ATAC_) as the input, the corresponding gene-expression data of that cell 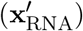 can be predicted as

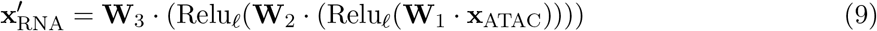

where **W**_*i*_ represents the weights of the *i*th layer. When learning the weights, we use the coarse GAM **A** (which is a binary matrix) as prior information to constrain the training procedure. We assume that **A** includes all the potential regulations between regions and genes, that is, the 0s in **A** are correct information but 1s in **A** can be false positives. Given this assumption, we construct a regularization term *L*_GAM_ to penalize the non-zero regulation strength from a region to a gene in the learned gene activity module that should be zero according to **A**:

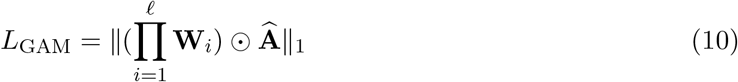

⊙ denotes the element-wise multiplication between two matrices and **Â** is the element-wise reversion of **A**. We use 𝓁_1_ norm to enforce the sparsity of the learned regulation strength.

After finishing the training procedure and having learned **W**_1_, **W**_2_ and **W**_3_, we then obtain a trained gene activity module which represents the complex nonlinear gene activity function between the scATAC-seq and the scRNA-seq data.

### Using anchor information

Root cell information is often needed when performing trajectory inference algorithm. Such information can also be utilized in `scDART` as “anchor”. That is, the root cells in the scRNA-seq dataset should be matched with the root cells in the scATAC-seq dataset. These cells are also called anchor cells. Other cells which are not root cells can also be anchor cells if we know their matching information across the two modalities. An additional loss term is added to the overall loss function when anchor cell information is used:

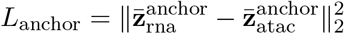

where 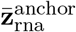 is the mean latent embedding of anchor cells within the scRNA-seq dataset, and 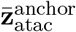 is the mean latent embedding of anchor cells within the scATAC-seq dataset. The loss make the anchor to match in the latent space by forcing the mean of the anchor cells’ latent distribution to be closer. The weight for this loss term is 1.

### Training scDART

When training `scDART`, the parameters in both the gene activity function module (**Θ**_proj_) and the projection module (**Θ**_gact_) are learned to minimize the overall loss function (Equ. 1). The training processes are different between scRNA-seq and scATAC-seq batches. When training with the scATAC-seq batch, we feed the data into the gene activity function module, and take the transformed pseudo-scRNA-seq data into the projection module to obtain **Z**_ATAC_. We use stochastic gradient descent to update parameters in both modules and minimize Equ. 11 (part of Equ. 1 that is relevant to scATAC-seq batch).

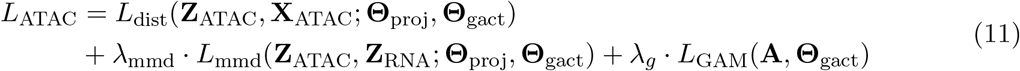

When training with scRNA-seq batch, we directly feed the data into the projection module to obtain **Z**_RNA_. We use stochastic gradient descent to update parameters in only the projection module and minimize Equ. 12 (part of Equ. 1 that is relevant to scRNA-seq batch).

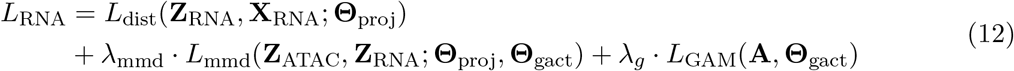

The training of `scDART-anchor` follows the same procedure as `scDART`. The only difference is that `scDART-anchor` includes the anchor loss *L*_anchor_(**Z**_ATAC_, **Z**_RNA_; **Θ**_proj_, **Θ**_gact_) in both Equ. 11 and Equ. 12 when training on scATAC-seq and scRNA-seq batches.

### Post-processing after training

After obtaining the latent embedding **Z**_ATAC_ and **Z**_RNA_, we apply a post-processing step to further refine the latent embedding to form a cleaner trajectory structure. We construct *k* (*k* = 10) mutual nearest neighbor graph [31] on the cells from **Z**_ATAC_ and **Z**_RNA_: for each cell in **Z**_RNA_, we find its *k* nearest cells in **Z**_ATAC_, and vice versa. After constructing the graph, we calculate weights on the graph. For cell *i* in **Z**_RNA_ and cell *j* in **Z**_ATAC_, the weight is:.

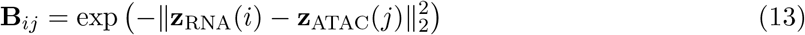

then we update the latent embedding of each cell by the embedding of its neighbors. For example, for each cell *i* in the scRNA-seq data, the new latent embedding 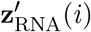 is calculated as:

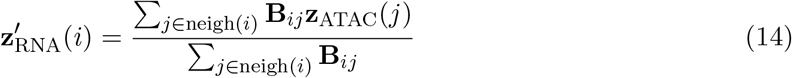

The post-processing step makes different trajectory branches more distinguishable in the latent space, thus helping trajectory inference methods to detect more accurate trajectories in complex trajectory structures.

### Trajectory inference on the integrated latent space

`scDART` outputs latent space representations of the integrated data, and then any trajectory inference algorithm that takes the reduced dimensional space representation can be used to infer the cell trajectories, such as diffusion pseudotime (DPT) [14], Slingshot [32], and Monocle [33]. In our tests, we apply DPT [14] on the latent embedding to infer the pseudotime for cells from both modalities jointly. Our backbone inference procedure is similar to the procedure used in PAGA [34]. When inferring the trajectory backbone from the latent embedding, we first run Leiden clustering [35] on the latent embedding, then construct a fully connected graph on the cluster centroids with the pairwise Euclidean distance between cluster centroids as the weights of the edges between them, and run minimum spanning tree to infer the trajectory backbone on the cluster centroids.

### Differential expression analysis

We find differentially expressed genes and accessible motifs along the trajectory by testing the significance of their changes depending on the pseudotime. We use likelihood ratio test as the significant test.

The alternative hypothesis assumes that the change of gene or motif depends on the pseudotime. We use a generalized additive model to fit their expression or accessibility values with pseudotime:

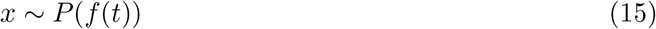

where *f* (*t*) is build with degree-4 spline functions. For the link function *P* (·), we assume that the log-transformed gene expression and motif follow Gaussian distribution:

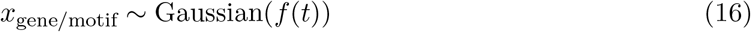

The null hypothesis assumes that:

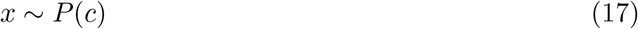

where *c* is a constant.

We then compare the two nested models using likelihood ratio test. We conduct the test for every gene and motif, and sort them separately according to their p-values. The significant genes and motifs are selected based on both their p-values and their relative ordering. We select the genes with p-values smaller than 0.05 and total number 100 cut-off and select the motifs with p-values smaller than 0.05 and total number 50 cut-off.

## Data simulation

The simulated scRNA-seq and scATAC-seq data are generated with an extended version of SymSim [36] which simulates scRNA-seq data. In SymSim, a kinetic model is used to model the mRNA counts in cells, where a gene is considered to be either in an *on* state or in an *off* state [37]. When a gene is in the *on* state, its transcripts are synthesized with rate *s*, and synthesized mRNAs degrade with a rate *d*. A parameter *k*_on_ represents the rate at which a gene enters the *on* state, and *k*_off_ represents the rate of the gene entering the *off* state. To generate multiple discrete or continuous cell types, SymSim defines an “identity vector” for each cell, and the identity vectors can evolve along a user-provided tree which represents the trajectory backbone.

In this work, we extended SymSim so that it also generates scATAC-seq data. Additional file 1: Figure S5a shows the process of generating *N* cells which have both scRNA-seq and scATAC-seq data. Denote the number of genes by *G* and the number of regions by *R*. A binary *R × G* GAM is provided to represent which regions affect which genes. As the scRNA-seq data depends on the scATAC-seq data, we first generate the scATAC-seq data. Similar to how SymSim generates scRNA-seq data along a continuous trajectory, we start with a “cell chromatin accessibility identity vector” of length *v* for the root cell and let it evolve along the given trajectory structure through a Brownian motion process to generate the “cell chromatin accessibility identity vectors” of cells along the tree. Each region has a “region identity vector” which is of the same length *v*. Multiplying the “cell chromatin accessibility identity matrix” and the “region identity vector matrix” we obtain an *N × R* matrix, where entries with larger values correspond to higher chromatin accessibility. We call this matrix “non-realistic scATAC-seq data” as its distribution is not the same as the distribution in real data. We then map the data in this matrix to a distribution obtained from a real scATAC-seq dataset [38] to get the realistic scATAC-seq data.

The scRNA-seq data is affected by both the input trajectory tree and the scATAC-seq data. We first generate the kinetic parameters for generating scRNA-seq data in the same way as in SymSim, and obtain the “realistic kinetic parameter matrix” shown in Additional file 1: Figure S5a. We now use the scATAC-seq data and the GAM to adjust *k*_on_, as we consider that the accessibility of the associated regions of a gene affects the rate that the gene is switched on, which is what *k*_on_ corresponds to. Now among the three kinetic parameters of scRNA-seq data, *k*_on_ is affected by scATAC-seq data, and *k*_off_ and *s* are affected by the input trajectory, thus we have combined both the effects of chromatin accessibility and cell differentiation process into the final scRNA-seq data. We then add technical noise to the scRNA-seq data, and divide all cells into two batches while adding batch effects. To mimic the unmatched data, for one batch we keep only the scRNA-seq data and for the other batch we keep only the scATAC-seq data.

### Evaluation metrics

When ground truth information is available, we use the following metrics to evaluate the latent embedding learned by `scDART` and the trajectories inferred based on it: neighborhood overlap score, F1 score [23] and Kendall-*τ* score [39] on inferred pseudotime. With simulated data, we also evaluate the gene activity module learned by `scDART`. Given scATAC-seq data of a cell, we use the gene activity module of `scDART` to generate its pseudo-scRNA-seq data, and measure the normalized mean square error (MSE) between pseudo-scRNA-seq data and the ground truth scRNA-seq data of the cell.

Neighborhood overlap score [12, 7] can be used to measure how well datasets are integrated when there exists cell-cell correspondence across data modalities. Given a neighborhood size *k*, it constructs a *k*-nearest neighbor graph on the latent embedding of cells from both scRNA-seq and scATAC-seq data, and calculates the proportion of cells that have their corresponding cells in the other modality included within its neighborhood.

The latent embedding of the integrated data is evaluated through both visualization and the quantitative accuracy of the inferred trajectories. The accuracy of trajectory is measured from two different aspects: the accuracy of cell branch assignment, and the accuracy of cell pseudotime assignment. We measure cell branch assignment using F1 score which was used in [23]. Here we briefly describe the calculation of F1 score. Given the ground truth and inferred cell branch assignment, we first calculate the Jaccard similarity between every pair of inferred and ground truth cell branches. For every two cell branch, the Jaccard similarity is calculated as the size of their intersection cell sets over the size of their union cell sets. For every branch in ground truth or inferred trajectory, we calculate its “maximum Jaccard similarity” as the maximum value out of its Jacaard similarities with all branches in the inferred/ground truth trajectory. Then we can calculate the *recovery* as the average maximum Jaccard similarity for every branch in ground truth and the *relevance* as the average maximum Jaccard similarity for every branch in inferred branches. The F1 score is then calculated as

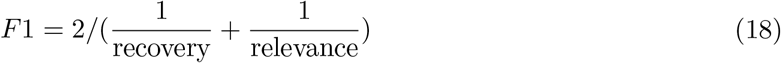

F1 score lies within the range between 0 and 1. The higher the score is, the better cell branches are assigned. We measure cell pseudotime assignment using Kendall-*τ* score [39], which is a rank-based correlation measurement that is commonly used to measure pseudotime inference accuracy [40, 32]. Kendall-*τ* score lies within the range between −1 and 1. A higher Kendall-*τ* score means a better pseudotime inference accuracy.

In simulated datasets, we can retrieve the ground truth gene expression data for cells in the scATAC-seq batch. Then we can measure how well the pseudo-scRNA-seq data learned from the gene activity module matches the ground truth scRNA-seq data using normalized MSE. Denoting the pseudo-scRNA-seq data of cell *i* as 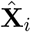, and the ground truth scRNA-seq data as **X**_*i*_, the normalized MSE is calculated as

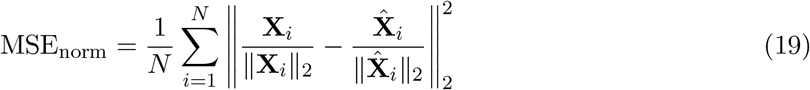

where *N* is the total number of cells.

## Ethics approval and consent to participate

Not applicable.

## Consent for publication

Not applicable.

## Availability of data and materials

scDART is implemented as both Python packages, which are freely available under the GPL-3 license. Source codes have been deposited at the GitHub repository (https://github.com/PeterZZQ/scDART). The testing script has been deposited at the GitHub repository (https://github.com/PeterZZQ/scDART_test). The datasets analyzed in this study are available from the Gene Expression Omnibus (GEO) repository under the following accession numbers: GSE126074 [1], GSE137117 [19], GSE96772 [21], and GSE117498 [22].

## Author’s contributions

ZZ and XZ conceived the project. ZZ, CY implement the model. XZ supervised the research. ZZ and XZ contributed to the writing of the manuscript. All authors read and approved the final manuscript.

## Acknowledgements

The authors would like to thank Dr. Xi Chen (from SUSTech) for helpful discussions.

## Supplementary Figures

**Figure S1:**
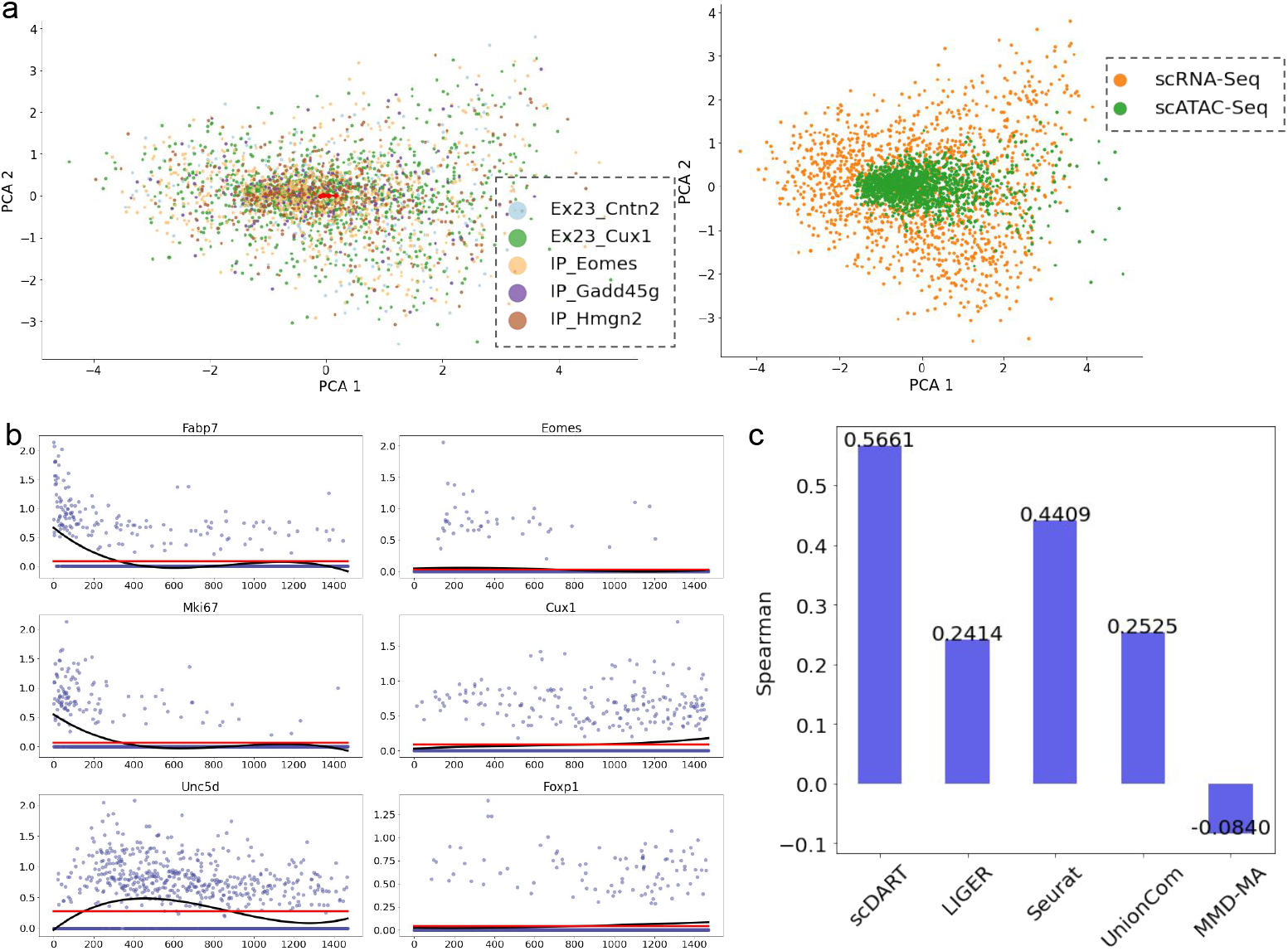
Additional test results on the SNARE-seq mouse neonatal brain cortex dataset. **a** Latent embedding of integrated data obtained with `MMD-MA`. Left: cells colored with cell type annotations from the original paper; Right: cells colored with batches (or modality). **b** Expression levels of *Mki67, Fabp7, Unc5d, Cux1, Eomes*, and *Foxp1* along the pseudotime inferred from measured scRNA-seq data. The black and red lines correspond to the fitted statistical models under alternative and null hypothesis, respectively, when conducting likelihood ratio test. **c** Pseudotime consistency score, where Spearman correlation is used.

**Figure S2:**
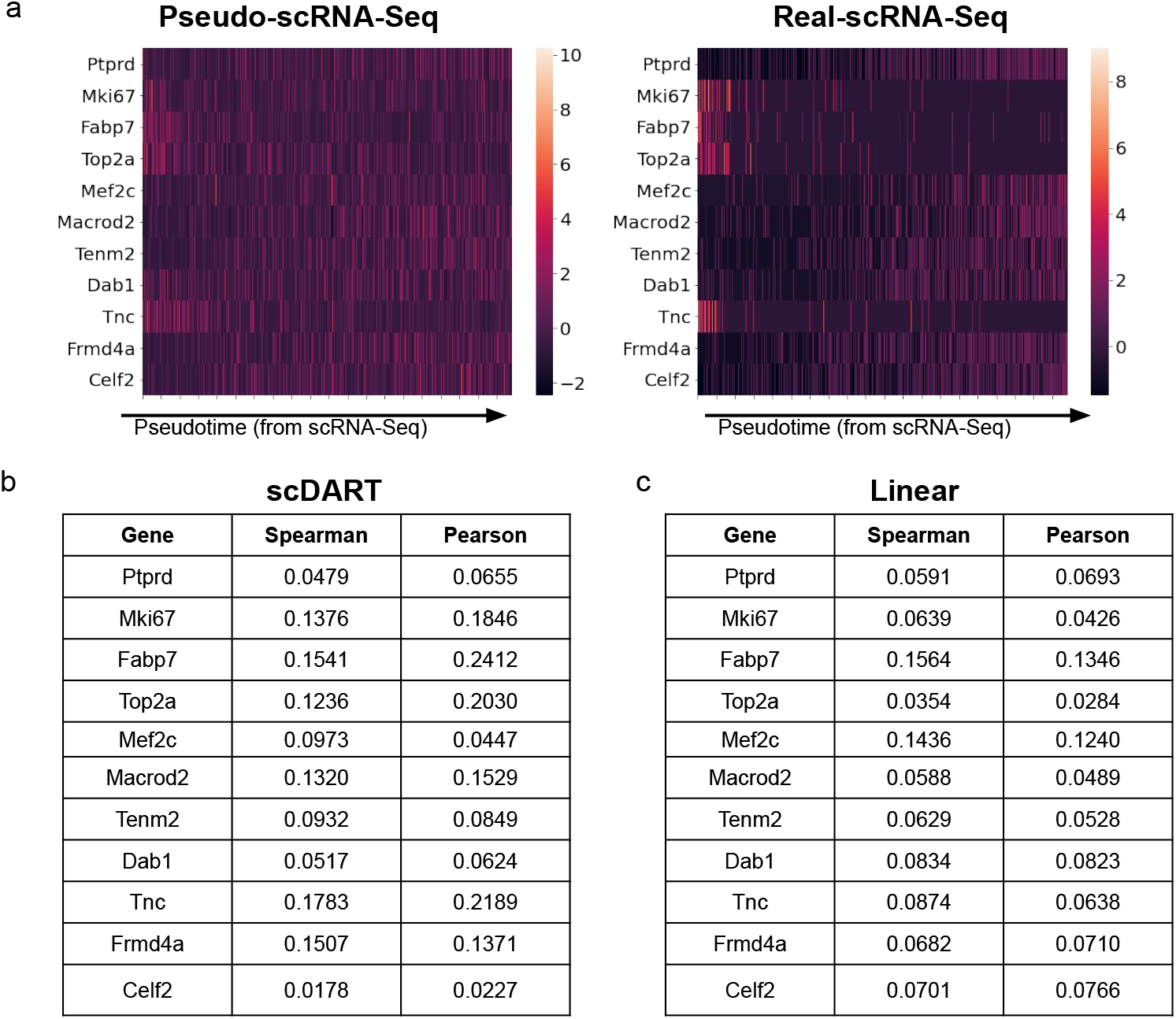
Comparison between pseudo-scRNA-seq and real-scRNA-seq. **a** Pseudo- (left) and real- scRNA-seq (right) data of selected DE genes in cells ordered on the x-axis according to pseudotime inferred from real scRNA-seq data. Colors show the gene expression levels normalized into z-scores. **b** Correlation scores between pseudo- and real scRNA-seq data of top DE genes. The pseudo-scRNA-seq data is obtained with `scDART`. **c** Correlation score between pseudo- and real-scRNA-seq data. The pseudo-scRNA-seq data is obtained with linear transformation on the scATAC-seq data with the input GAM.

**Figure S3:**
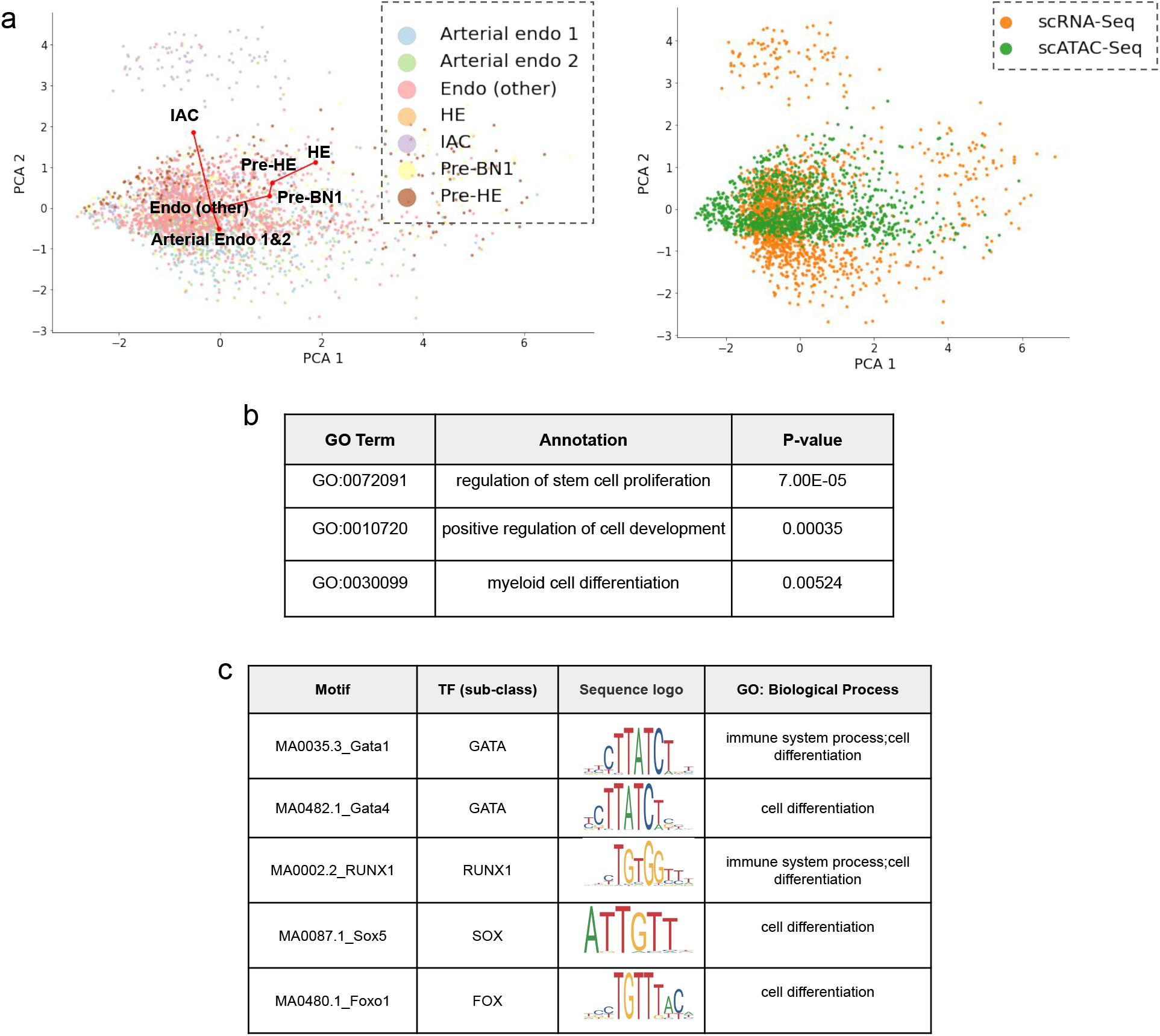
Additional test results on the mouse endothelial dataset. **a** The latent embedding of `MMD-MA` on mouse endothelial dataset. Left: cells colored with cell type annotations from the original paper, and the red lines show the inferred trajectory backbone; Right: cells colored with batches (or modality). **b** Top gene ontology terms of DE genes on mouse endothelial cell development dataset. **c** Selected differentially accessible motifs on mouse endothelial cell development dataset.

**Figure S4:**
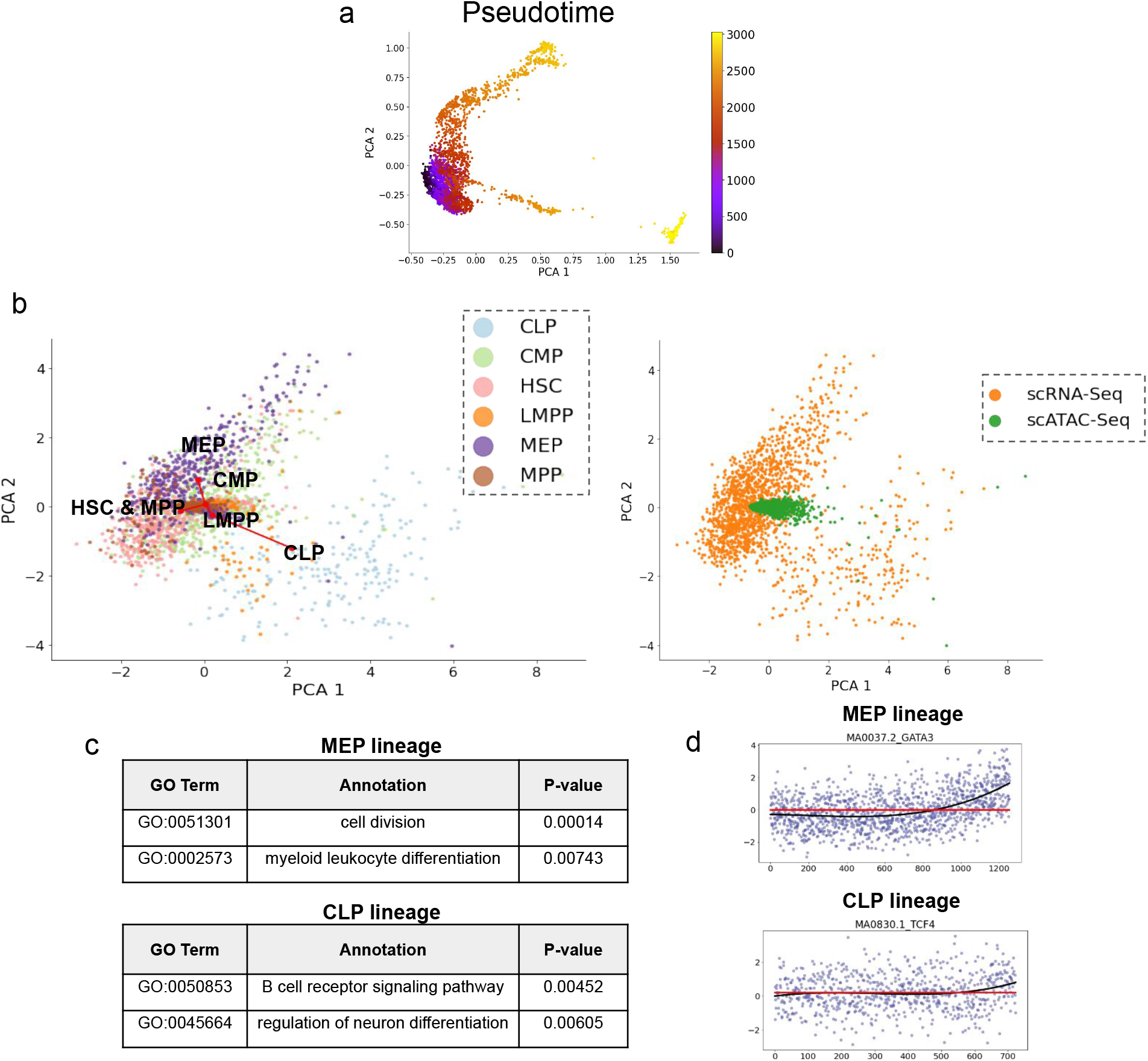
Additional test results on the human hematopoiesis dataset. **a** Latent embedding of `scDART` on human hematopoiesis dataset, where cells are colored with the inferred pseudotime. **b** The latent embedding of `MMD-MA` on human hematopoiesis dataset. Left: cells colored with cell type annotations from the original paper, and the red lines show the inferred trajectory backbone; Right: cells colored with batches (or modality). **c** The top gene ontology terms found by `TopGO` on human hematopoiesis dataset. **d** The deviation value (from ChromVAR) of differentially accessible motifs along MEP and CLP lineages. The black and red lines correspond to the fitted statistical models under alternative and null hypothesis, respectively, when conducting likelihood ratio test.

**Figure S5:**
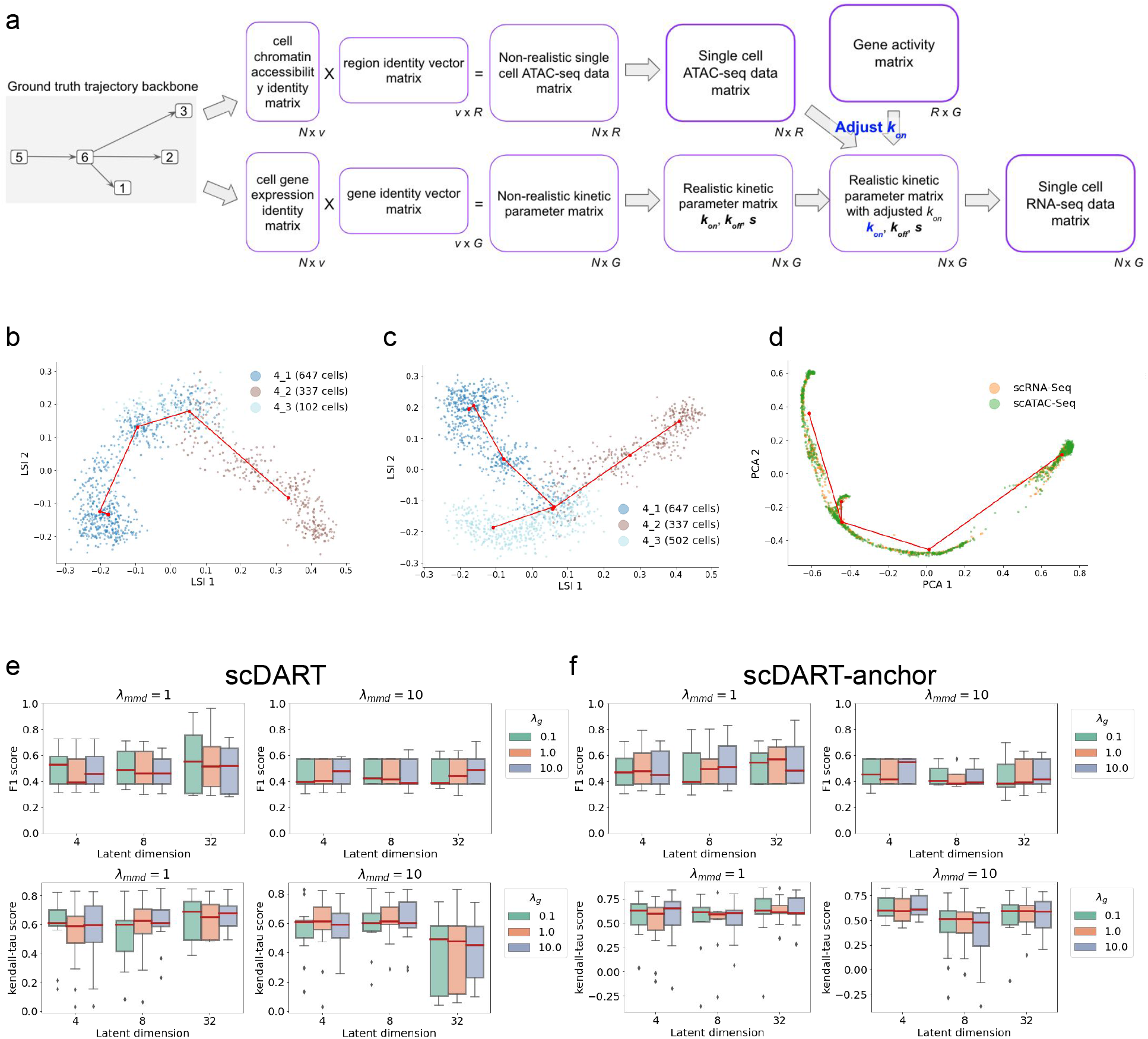
Additional test results on simulated datasets. **a** Illustration of the data simulation process. Given the ground truth trajectory and the gene activity matrix, the simulation procedure generates both scRNA-seq and scATAC-seq data. **b** The trajectory backbone learned from scATAC-seq where branch 4 3 has only 102 cells. **c** The trajectory backbone learned from scATAC-seq where branch 4 3 has 502 cells. **d** The trajectory backbone learned from `scDART` latent embedding where cells are colored by data batches. **e-f**. The boxplots of F1-score and Kendall-*τ* score under different hyper-parameter settings of (e) `scDART` and (f) `scDART-anchor`.

